# Discovery of physiological and cancer-related regulators of 3’ UTR processing with KAPAC

**DOI:** 10.1101/195958

**Authors:** Andreas J. Gruber, Ralf Schmidt, Souvik Ghosh, Georges Martin, Andreas R. Gruber, Erik van Nimwegen, Mihaela Zavolan

## Abstract

3’ UTR length is regulated in relation to cellular state. To uncover key regulators of poly(A) site (PAS) use in specific conditions, we have developed PAQR, a method for quantifying PAS use from RNA sequencing data and KAPAC, an approach that infers activities of oligomeric sequence motifs on PAS choice. Application of PAQR and KAPAC to RNA sequencing data from normal and tumor tissue samples uncovered sequence motifs that can explain changes in cleavage and polyadenylation in specific cancers. In particular, our analysis points to Polypyrimidine tract binding protein 1 as a regulator of PAS choice in glioblastoma.

## Background

The 3’ ends of most eukaryotic mRNAs are generated through endonucleolytic cleavage and polyadenylation (CPA) [1–3]. These steps are carried out in mammalian cells by a 3’ end processing complex composed of the cleavage and polyadenylation specificity factor (which includes the proteins CPSF1 (also known as CPSF160), CPSF2 (CPSF100), CPSF3 (CPSF73), CPSF4 (CPSF30), FIP1L1 and WDR33), the mammalian cleavage factor I (CFIm, a tetramer of two small, NUDT21 (CFIm 25) subunits, and two large subunits, of CPSF7 (CFIm 59) and/or CPSF6 (CFIm 68)), the cleavage factor II (composed of CLP1 and PCF11), the cleavage stimulation factor (CstF, a trimer of CSTF1 (CstF50), CSTF2 (Cstf64) and CSTF3 (CstF77)), symplekin (SYMPK), the poly(A) polymerase (PAPOLA, PAPOLB, PAPOLG) and the nuclear poly(A) binding protein (PABPN1) [3, 4], Crosslinking and immunoprecipitation (CLIP) revealed the distribution of core 3’ end processing factor binding sites in pre-mRNAs [5] and the minimal polyadenylation specificity factor that recognizes the polyadenylation signal, consisting of the CPSF1, CPSF4, FIP1L1, and WDR33 proteins, has been identified [6, 7].

Most genes have multiple poly(A) sites (PAS), which are differentially processed across cell types [8], likely due to cell type-specific interactions with RBPs. The length of 3’ UTRs is most strongly dependent on the mammalian cleavage factor I (CFIm), which promotes the use of distal poly(A) sites [5, 9–12], Reduced expression of CFIm 25 has been linked to 3’ UTR shortening, cell proliferation and oncogene expression in glioblastoma cell lines [11], while increased levels of CFIm 25 due to gene duplication has been linked to intellectual disability [13], The CSTF2 component of the CstF subcomplex also contributes to the selection of poly(A) sites [5, 14], but in contrast to CFIm, depletion of CSTF2 leads to increased use of distal poly(A) sites (dPAS), especially when the paralogous CSTF2T is also depleted [14], PCF11 and FIP1L1 proteins similarly promote the use of proximal poly(A) sites (pPAS) [12].

Many splicing factors modulate 3’ end processing. Most strikingly, the U1 small nuclear ribonucleoprotein (snRNP) promotes transcription, masking poly(A) sites whose processing would lead to premature CPA, through a ‘telescripting’ mechanism [15, 16], The U2AF65 spliceosomal protein interacts with CFIm [17] and competes directly with the heterogeneous nucleoprotein C (HNRNPC) for binding to uridine(U)-rich elements, regulating the splicing and thereby exonization of Alu elements [18], HNRNPC represses CPA at poly(A) sites where U-rich sequence motifs occur [19], Other splicing factors that have been linked to poly(A) site selection are the neuron-specific NOVA1 protein [20], the nuclear and cytoplasmic poly(A) binding proteins [12, 21], the heterogeneous ribonucleoprotein K (HNRNPK) [22], and the poly(C) binding protein (PCBP1) [23], However, the mechanisms remain poorly understood. An emerging paradigm is that position-dependent interactions of pre-mRNAs with RBPs influence poly(A) site selection, as well as splicing [24], By combining mapping of RBP binding sites with measurements of isoform expression, Ule and colleagues started to construct ‘RNA maps’ relating the position of cis-acting elements to the processing of individual exons [25], However, whether the impact of a regulator can be inferred solely from RNA sequencing data obtained from samples with different expression levels of various regulators is not known.

To address this problem, we have developed KAPAC (for **k**-mer **a**ctivity on **p**oly**a**denylation site **c**hoice), a method that infers position-dependent activities of sequence motifs on 3’ end processing from changes in poly(A) site usage between conditions. By analogy with ‘RNA maps’, and to emphasize the fact that our approach does not use information about RBP binding to RNA targets, we summarize the activities of individual motifs inferred by KAPAC from different regions relative to poly(A) sites as ‘impact maps’. As 3’ end sequencing remains relatively uncommon, we have also developed PAQR, a method for **p**oly**a**denylation site usage **q**uantification from **R**NA sequencing data, that allows us to evaluate 3’ end processing in data sets such as those from The Cancer Genome Atlas (TCGA) Research Network [26], We demonstrate that KAPAC identifies binding motifs and position-dependent activities of regulators of CPA from RNA-seq data obtained upon the knock-down of these RBPs, and in particular, that CFIm promotes CPA at poly(A) sites located ∼50 to 100 nucleotides (nt) downstream of the CFIm binding motifs. KAPAC analysis of TCGA data reveals pyrimidine-rich elements associated with the use of poly(A) sites in cancer and implicates the polypyrimidine tract-binding protein 1 (PTBP1) in the regulation of 3’ end processing in glioblastoma.

## Results

### Inferring sequence motifs active on PAS selection with KAPAC

As binding specificities of RBPs have only recently been started to be determined *in vivo* in high-throughput [27], we developed an unbiased approach, evaluating the activity of all possible sequences of length *k* (*k*-mers, with *k* in the range of RBP-binding site length, 3-6 nucleotides [28]) on PAS usage. Briefly, we first compute the relative use of each PAS *p* among the *P* poly(A) sites (*P* > 1) in a given terminal exon across all samples *s*, as 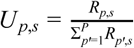, where *R_p,s_* is the number of reads observed for poly(A) site *p* in sample *s* (Figure 1A). KAPAC aims to explain the observed changes in relative poly(A) site usage *U_p,s_* in terms of the activity of a *k*-mer *k* within a sample *s* and the excess counts (over the background expected based on the mononucleotide frequencies, see section 2.2.1 of the Supplementary methods) *N_p,k_* of the *k*-mer within a region located at a specific distance relative to the poly(A) site *p* (Figure 1B-C). Running KAPAC for regions located at various relative distances with respect to the PAS (Figure 1D) allows the identification of the most significantly active *k*-mers as well as their location.

**Figure 1:**
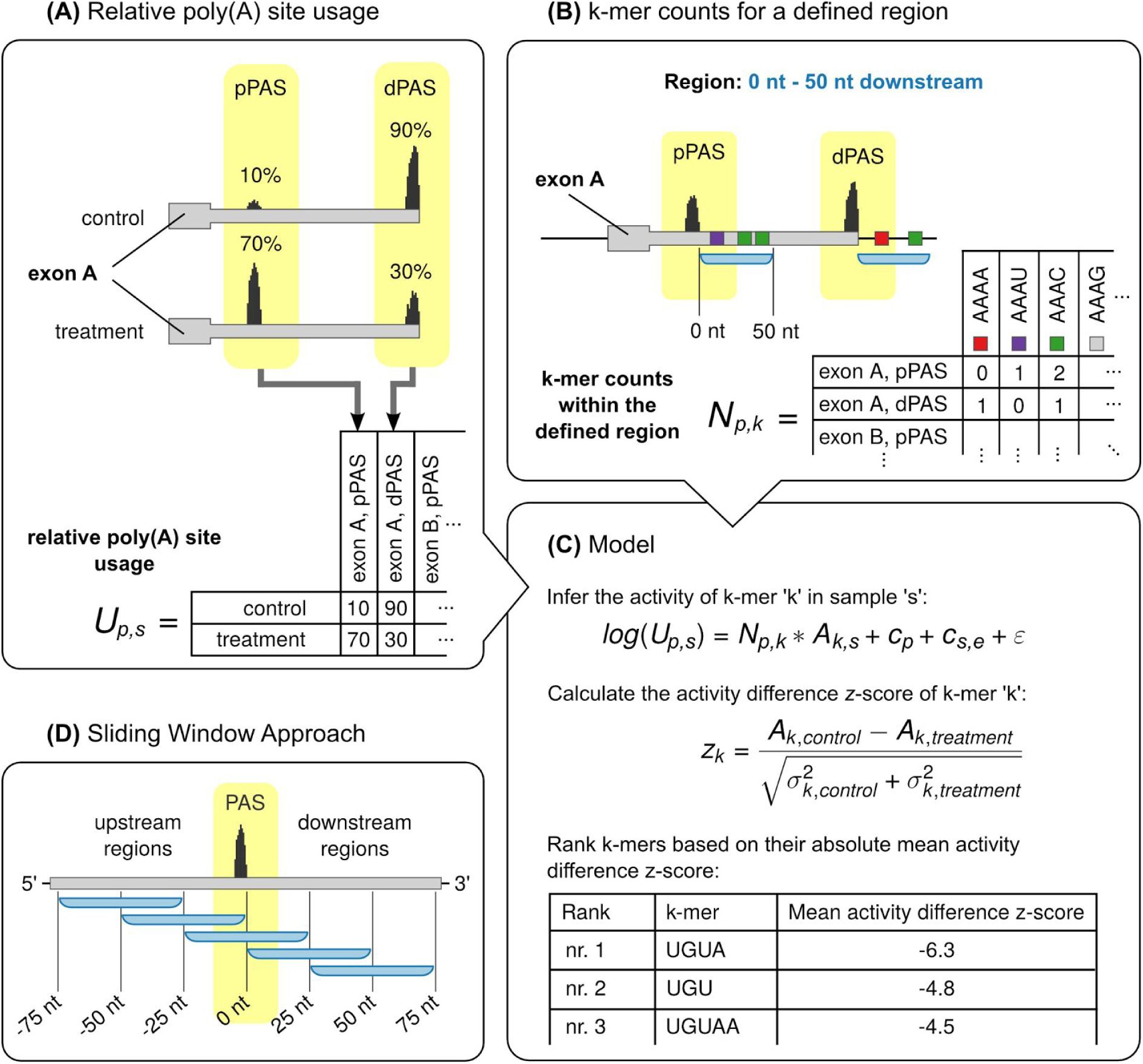
Schematic outline of the KAPAC approach. **(A)** Tabulation of the relative usage of poly(A) sites in different experimental conditions (here: control and treatment). **(B)** Tabulation of *k*-mer counts for regions (blue) located at a defined distances with respect to poly(A) sites *p*. **(C)** Based on the usage of poly(A) sites relative to the mean across samples and the counts of *k*-mers *k* in windows located at specific distances from the poly(A) sites *p*, KAPAC infers activities *A_k,s_* of *k*-mers in samples *s*. *c_s,e_* is the mean relative usage of poly(A) sites from exon *e* in sample *s*, *c_p_* is the mean log2-relative usage of poly(A) site *p* across samples, and *ε* is the residual error. KAPAC ranks *k*-mers based on the absolute z-score of the mean activity difference in two conditions (here: in control relative to treatment). **(D)** Fitting the KAPAC model for windows located at specific distances relative to poly(A) sites, position-dependent activities of sequence motifs on poly(A) site use are inferred.

### KAPAC uncovers expected position-specific activities of RBPs on pre-mRNA 3’ end processing

To evaluate KAPAC we first analyzed PAS usage data obtained by 3’ end sequencing upon perturbation of known RBP regulators of CPA. Consistent with the initial study of poly(C) binding protein 1 (PCBP1) role in CPA [23], as well as with the density of its CCC - (C)_3_ - binding element around PAS that do and PAS that do not respond to PCBP1 knock-down (Figure 2A), KAPAC revealed that (C)_3_ motifs strongly activate the processing of poly(A) sites located 25-100 nucleotides downstream (Figures 2B-C, Supplementary Table 1).

**Figure 2:**
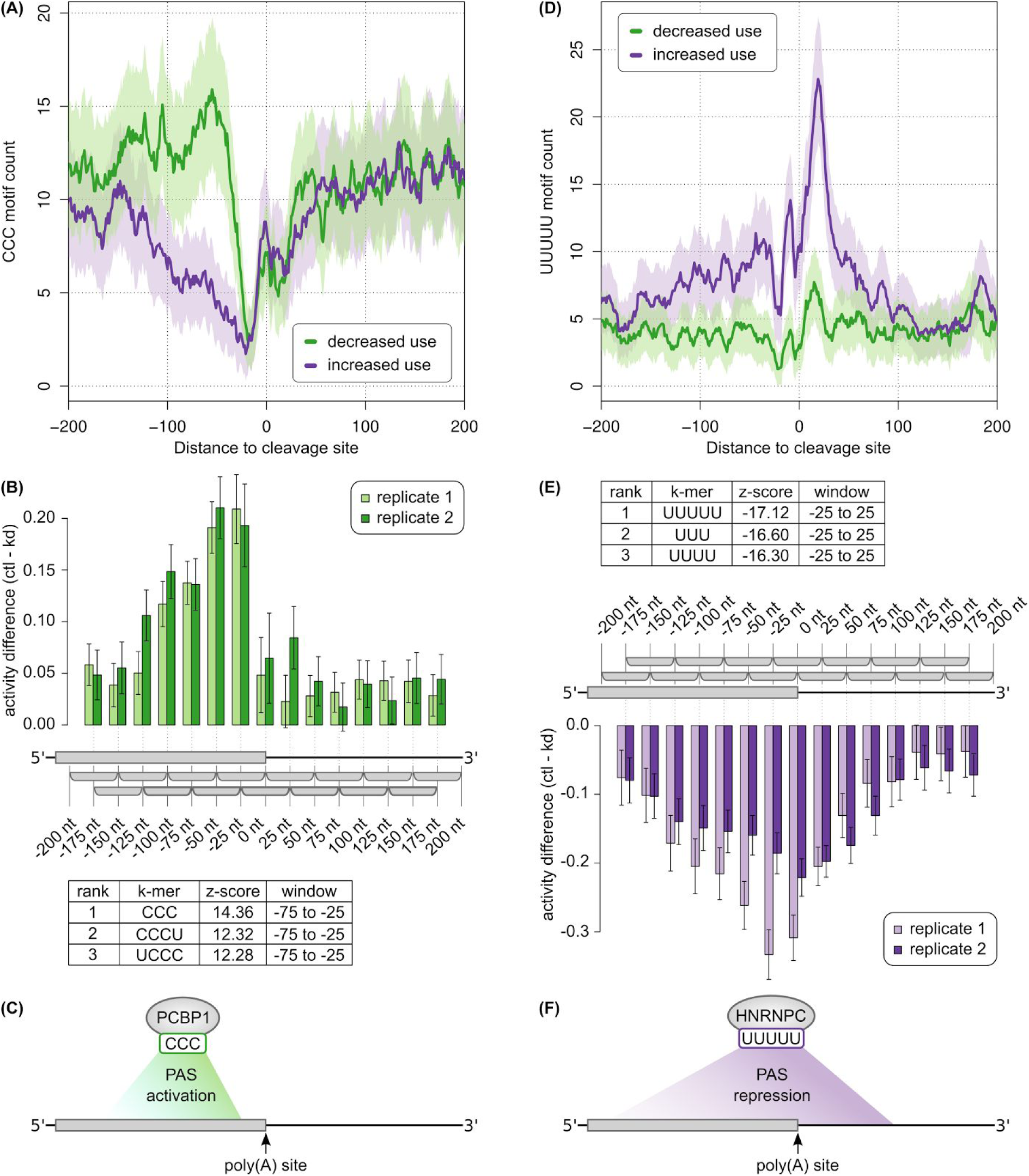
KAPAC accurately uncovers the activity of known regulators of poly(A) site choice. **(A)** Smoothened (+/-5nt) density of non-overlapping (C)_3_ motifs in the vicinity of poly(A) sites that are consistently processed (increased or decreased use) in two PCBP1 knock-down experiments from which 3’ end sequencing data is available [23], Shaded areas indicated standard deviations based on binomial sampling. **(B)** Difference of (C)_3_ motif activity inferred by KAPAC in the two replicates of control (Ctrl) versus PCBP1 knock-down (KD) experiments (number of PAS n = 3737). The positive differences indicate that (C)_3_ motifs are associated with increased PAS use in control samples. The table shows the three most significant motifs, with the z-score and position of the window from which they were inferred. **(C)** Model of the KAPAC-inferred impact of PCBP1 on CPA. **(D)** Smoothened (+/-5nt) density of non-overlapping (U)_5_ tracts in the vicinity of sites that are consistently processed (increased or decreased use) in two HNRNPC knock-down experiments [29], **(E)** Difference of (U)_5_ motif activity inferred by KAPAC in the two replicates of control (Ctrl) versus HNRNPC knock-down (KD) experiments (n = 4703). The negative differences indicate that (U)_5_ motifs are associated with decreased PAS use in the control samples. The table with the three most significant motifs is also shown, as in panel (B). **(F)** Model of the KAPAC-inferred impact of HNRNPC on CPA.

As in a previous study we found that the multi-functional heterogeneous ribonucleoprotein C (HNRNPC) modulates 3’ end processing (see also Figure 2D), we also applied KAPAC to 3’ end sequencing data obtained upon the knock-down of this protein. Indeed, we found that (U)_n_ sequences (*n* = 3 – 5 nucleotides) have a strongly repressive activity on poly(A) site choice, which, reminiscent of HNRNPC’s effect on exon inclusion [18], extends to a broad window, from approximately -200 nucleotides upstream to about 50 nucleotides downstream of poly(A) sites (Figure 2E-F, Supplementary Table 1). In contrast to the density of (U)_5_ motifs, which peaks immediately downstream of poly(A) sites, KAPAC inferred an equally high repressive activity of (U)_5_ motifs located upstream of the poly(A) site.

These results demonstrate that being provided only with estimates of poly(A) site expression in different conditions, KAPAC uncovers both the sequence specificity of the RBP whose expression was perturbed in the experiment, and the position-dependent, activating or repressing activity of the RBP on poly(A) site choice.

### The PAQR method to estimate relative PAS use from RNA-seq data

As 3’ end sequencing data remain relatively uncommon, we sought to quantify poly(A) site use from RNA sequencing data. The drop in coverage downstream of proximal PAS has been interpreted as evidence of PAS processing, generalized by the DaPars method to identify changes in 3’ end processing genome-wide [11], However, DaPars (with default settings) reported only 8 targets from the RNA-seq data obtained upon the knock-down of HNRNPC [29], and they did not include the previously validated HNRNPC target CD47 [19], whose distal PAS shows increased use upon HNRNPC knock-down (Figure 3A). Furthermore, DaPars quantifications of relative PAS use in replicate samples had limited reproducibility (Supplementary Figures 1, 2), as did the motif activities inferred by KAPAC based on these estimates (Figure 3B, Supplementary Figure 2). These results prompted us to develop PAQR, a method to quantify PAS use from RNA-seq data (Figure 3C). PAQR uses read coverage profiles to progressively segment 3’ UTRs at annotated poly(A) sites. At each step, it infers the breakpoint that decreases most the squared deviation from the mean coverage of a 3’ UTR segment when dividing the segment in two regions with distinct mean coverage (Figure 3C and Methods) relative to considering it as a single segment with one mean coverage. A key aspect of PAQR is that it only attempts to segment the 3’ UTRs at experimentally identified poly(A) sites, from an extensive catalog that was recently constructed [19], Using the HNRNPC knock-down data set that was obtained independently [29] for benchmarking, we found that the PAQR-based quantification of PAS use led to much more reproducible HNRNPC binding motif activity and more significant difference of mean z-scores between conditions (-22.92 with PAQR-based quantification vs. -10.19 with DaPars quantification, Figure 3B,D, Supplementary Figure 2). These results indicate that PAQR more accurately and reproducibly quantifies poly(A) site use from RNA-seq data.

**Figure 3:**
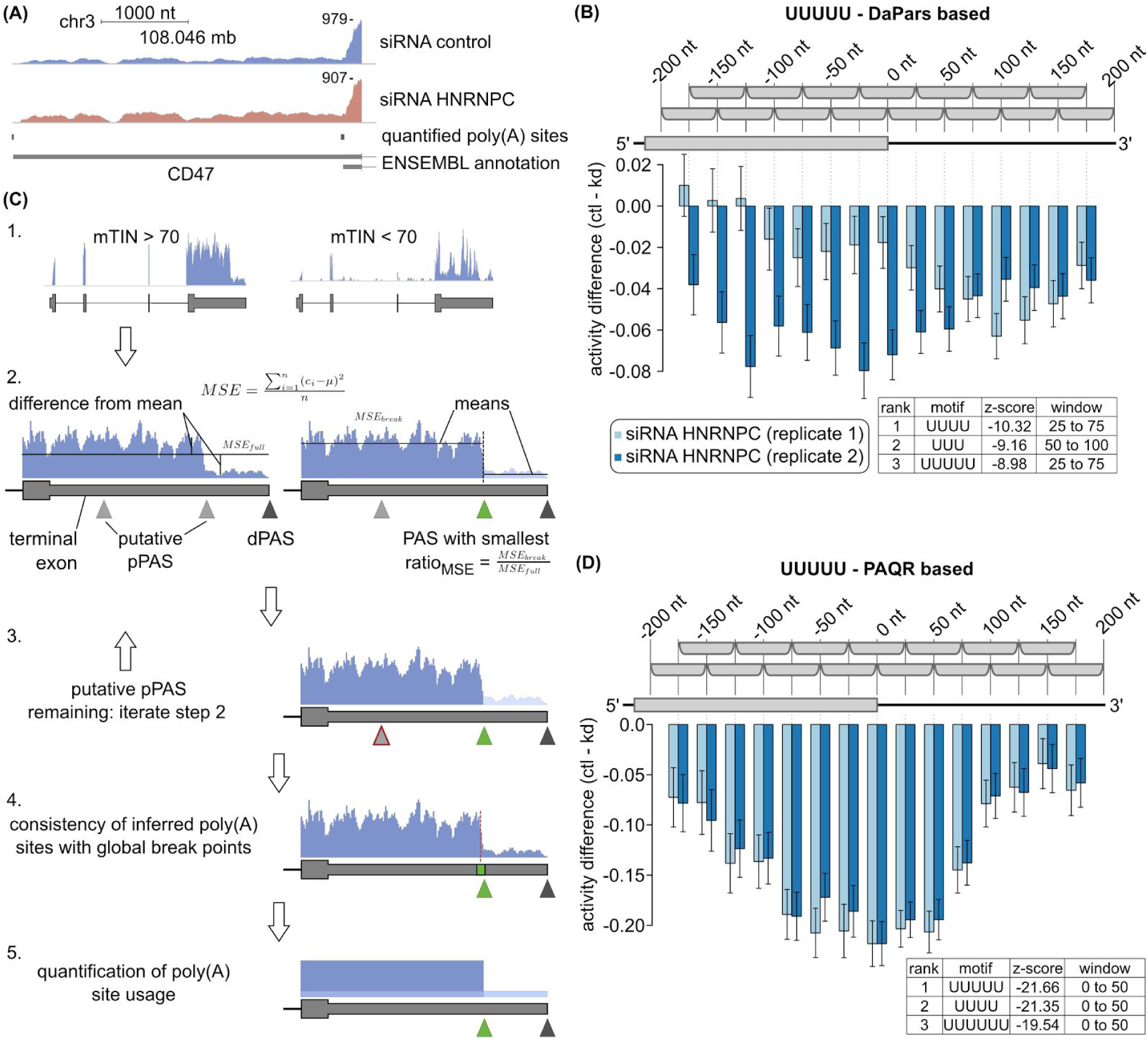
Overview on PAQR. **(A)** Read coverage profile of the CD47 terminal exon, whose processing is affected by the knock-down of HNRNPC [19], **(B)** KAPAC-inferred position-dependent activities of the (U)_5_ motif based on DaPars-based estimates of relative PAS use (number of PAS n = 13388) in the same data set as in (A). **(C)** Sketch of PAQR: (1.) Samples with highly biased read coverage along transcripts (low mTIN score), presumably affected by RNA degradation, are identified and excluded from the analysis; (2.) usage of proximal PAS (pPAS) in a sample is determined based on the expected drop in coverage downstream of the used PAS (ration of the mean squared deviation from mean coverage (MSE) in the full region compared to two distinct regions, split by the poly(A) site; (3.) step (2.) is repeated iteratively for subregions bounded by already determined PAS; (4.) the consistency between PAS called as used and the global best break points in corresponding regions is evaluated and in case of discrepancy, terminal exons are discarded from the analysis (5.) relative PAS use is calculated from the average read coverage of individual 3’ UTR segments, each corresponding to the terminal region of an isoform that ends at a used poly(A) site. **(D)** Similar HNRNPC activity on PAS use is inferred by KAPAC from estimates of PAS use generated either by PAQR from RNA sequencing data (n = 3599), or measured directly by 3’ end sequencing (Figure 2E).

### KAPAC reveals a position-dependent activity of CFIm binding on cleavage and polyadenylation

As KAPAC allows us to infer position-dependent effects of RBP binding on 3’ end processing, we next sought to unravel the mechanism of CFIm, the 3’ end processing factor with a relatively large impact on 3’ UTR length [5, 9, 10, 12], We thus depleted either the CFIm 25 or the CFIm 68 component of the CFIm complex by siRNA-mediated knock-down in HeLa cells, and carried out RNA 3’ end sequencing. As expected, CFIm depletion led to marked and reproducible 3’ UTR shortening (Figure 4A, see Methods for details). We found that the UGUA CFIm binding motif occurred with high frequency upstream of the distal poly(A) sites whose usage decreased upon CFIm knock-down, whereas it was rare in the vicinity of all other types of PAS (Figure 4B, Supplementary Table 1). These results indicate that CFIm promotes the processing of poly(A) sites that are located distally in 3’ UTRs and are strongly enriched in CFIm binding motifs in a broad region upstream of the poly(A) signal. KAPAC analysis supported this conclusion, further uncovering UGUA as the second most predictive motif for the changes in poly(A) site use in these experiments, after the canonical poly(A) signal AAUAAA (Figure 4C), which is also enriched at distal PAS [5], Interestingly, the activity profile further suggests that UGUA motifs located downstream of PAS may repress processing of these sites, leading to an apparent decreased motif activity when CFIm expression is high.

**Figure 4:**
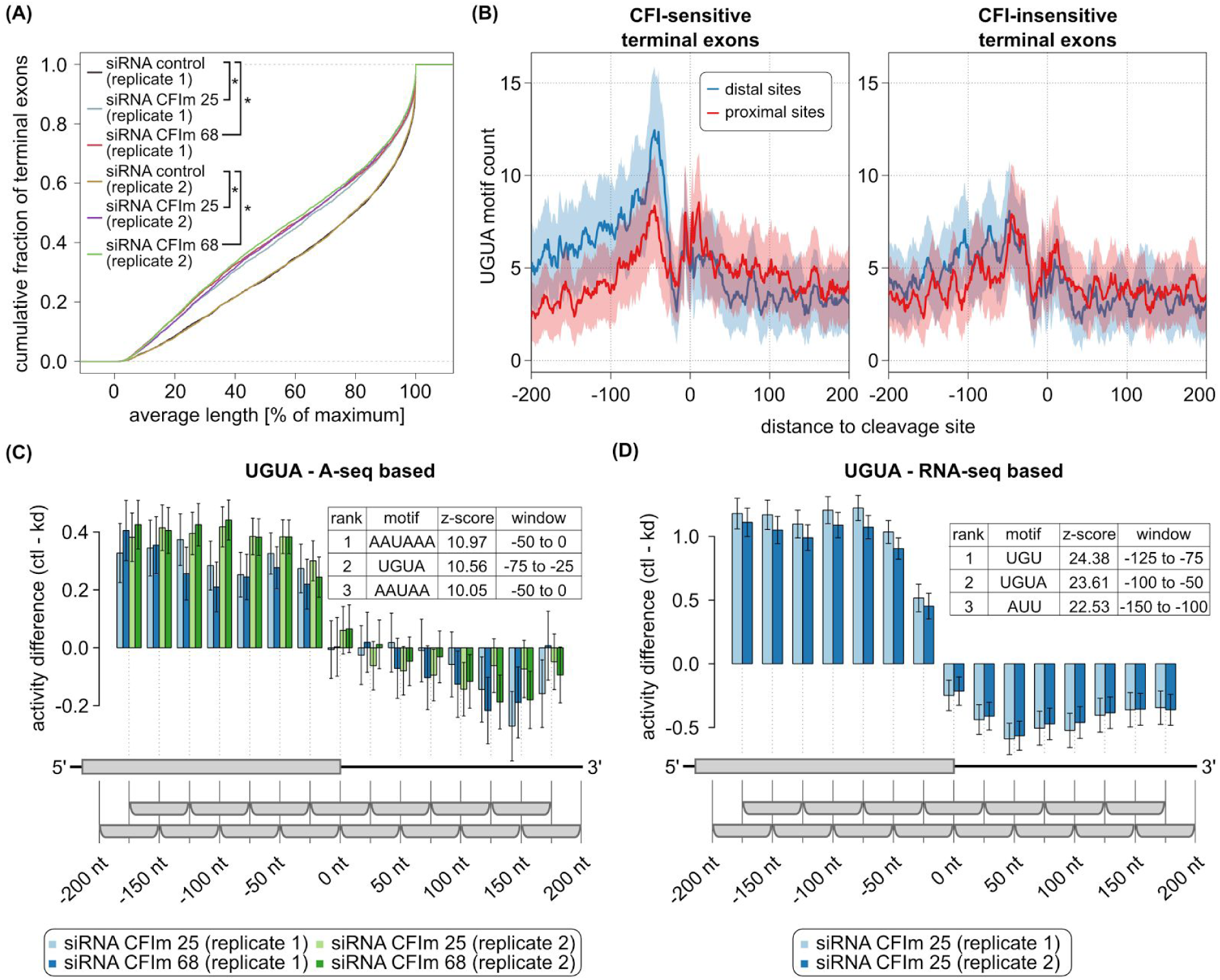
Position-dependent activation of pre-mRNA processing by CFIm. **(A)** The distributions of average terminal exon lengths (see Methods) computed from 5123 multi-PAS terminal exons quantified in CFIm 25, CFIm 68 knock-down and control samples indicate significant shortening of 3’ UTRs upon CFIm depletion (* two-sided wilcoxon signed-rank test p-value < 0.0001). **(B)** Smoothened (+/-5nt) UGUA motif density around PAS of terminal exons with exactly two quantified poly(A) sites, grouped according to the log fold change of the proximal-distal ratio (p/d ratio) upon CFIm knock-down. The left panel shows the UGUA motif frequency around the proximal and distal PAS of the 750 exons with the largest change in p/d ratio, while the right panel shows similar profiles for the 750 exons with the smallest change in p/d ratio. **(C)** KAPAC analysis of CFIm knock-down and control samples uncovers the poly(A) signal and UGUA motif as most significantly associated with changes in PAS usage (n = 3727). **(D)** UGUA motif activity is similar when the PAS quantification is done by PAQR from RNA sequencing data of CFIm 25 knock-down and control cells (n = 4287) [11].

We repeated these analyses on RNA-seq data obtained independently from HeLa cells depleted of CFIm 25 [11], obtaining similar activity profile (Figure 4D, Supplementary Table 2), including the apparent negative activity of sites that are located downstream on PAS processing. These results demonstrate that CFIm binds upstream of distal PAS to promote their usage, whereas binding of CFIm downstream of PAS may, in fact, inhibit processing of poly(A) sites.

### KAPAC implicates the pyrimidine tract binding proteins in 3’ end processing in glioblastoma

We then asked whether KAPAC can uncover a role of CFIm 25 in 3’ UTR shortening in glioblastoma (GBM), as has been previously suggested [11], We found that while 3’ UTRs are indeed markedly shortened in these tumors (Figure 5A), UGUA was not among the 20 motifs that most significantly explained the change in PAS usage in these samples. This may not be unexpected because in fact, once a certain threshold of RNA integrity is met, normal and tumor samples have CFIm expression in the same range (Supplementary Figure 5). Rather, KAPAC revealed that variants of the CU dinucleotide repeat, located from ~ 25 nt upstream to ~ 75 nt downstream of PAS, are most significantly associated with the change in PAS usage in tumors compared to normal samples (Figure 5B, Supplementary Table 3). Among the many proteins that can bind polypyrimidine motifs, the mRNA level of the pyrimidine tract binding protein 1 (PTBP1) was strongly anti-correlated with the median average length of terminal exons in this set of samples (Figure 5C). This suggested that PTBP1 masks the distally-located, CU repeat-containing PAS, which are processed only when PTBP1 expression is low, as it is in normal cells. Strikingly, KAPAC analysis of mRNA sequencing data obtained upon the double knock-down of PTBP1 and PTBP2 in HEK 293 cells (Supplementary Figure 3) [30] confirmed this hypothesis (Figure 5D). 181 of the 203 sites where the CU repeat motif was predicted to be active, were located most distally in the corresponding terminal exons. The PTBP1 crosslinking and immunoprecipitation data recently generated by the ENCODE consortium [31] confirmed the enriched binding of the protein downstream of the PAS of CU-containing, KAPAC-predicted target PAS (Figure 5E), whose relative usage decreases in tumor compared to control samples (Supplementary Figure 4). Furthermore, the enrichment of PTBP1-eCLIP reads is highest for the highest scoring PTBP1 targets (Figure 5F). These results more strongly implicate PTBP1 than CFIm 25 in the regulation of PAS use in glioblastoma.

**Figure 5:**
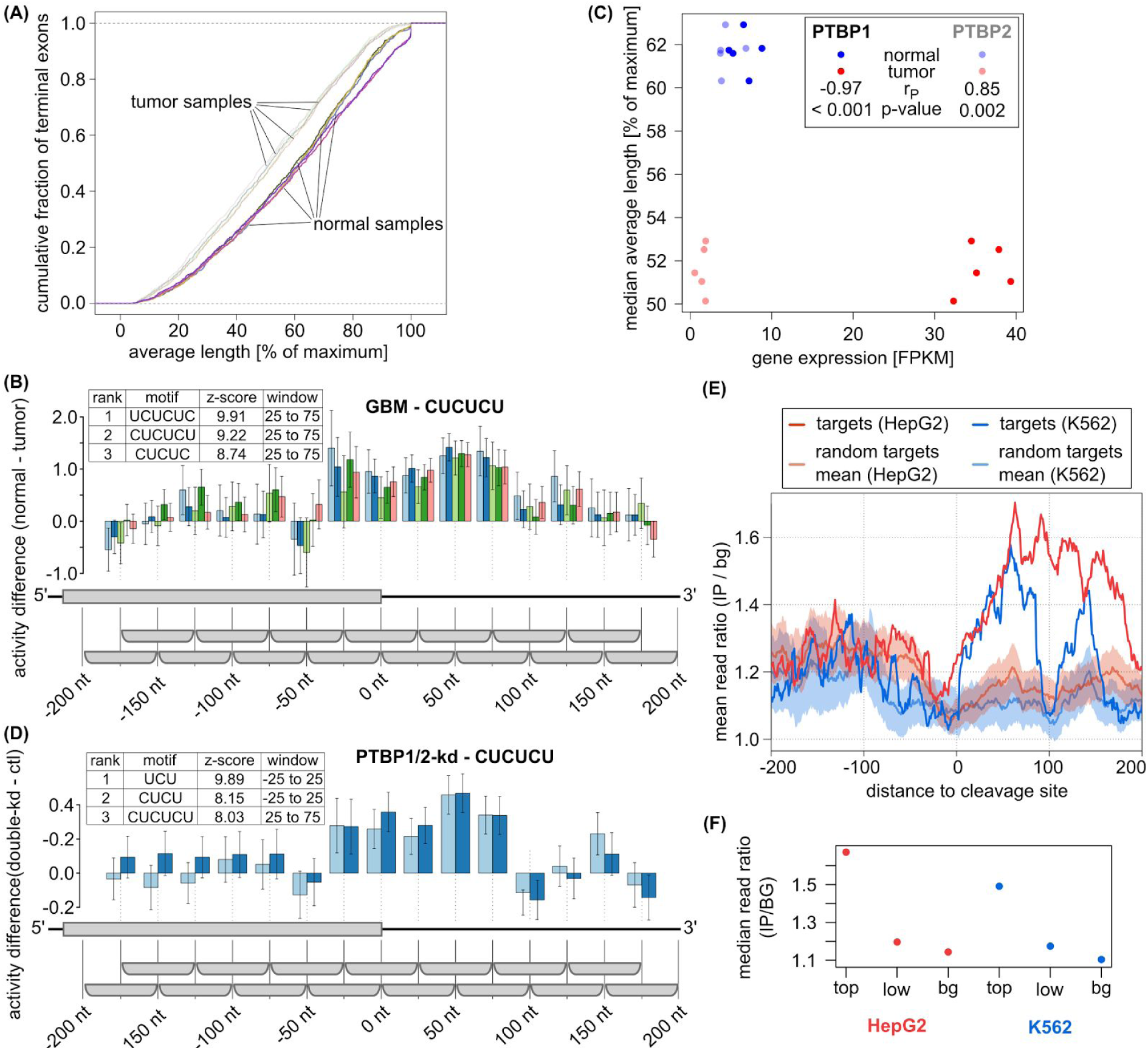
Regulation of PAS choice in glioblastoma samples from TCGA. **(A)** Cumulative distributions of weighted average length of 1172 terminal exons inferred by applying PAQR to five normal and five tumor samples (see Methods for the selection of these samples) show that terminal exons are significantly shortened in tumors. **(B)** Activity profile of CUCUCU, the second most significant motif associated with 3’ end processing changes in glioblastoma (number of PAS used in the inference n = 2119). The presence of the motif in a window from -25 to +75 relative to PAS is associated with increased processing of the site in normal tissue samples. **(C)** Expression of PTBP1 in the 10 samples from (A) is strongly anti-correlated (dark colored points, Pearson’s r (r_P_) = -0.97, p-value < 0.0001) with the median average length of terminal exons in these samples. In contrast, the expression of PTBP2 changes little in tumors compared to normal tissue samples, and has a positive correlation with terminal exon length (light colored points, r_P_ = 0.85, p-value = 0.002). **(D)** Activity profile of the same CUCUCU motif in the PTBP1/2 double knock-down (where the motif ranked third) compared to control samples (two biological replicates from HEK cells, number of PAS n = 2493). **(E)** Position-dependent PTBP1 binding inferred from two eCLIP studies (in HepG2 (thick red line) and K562 (thick blue line) cell lines) by the ENCODE consortium is significantly enriched downstream of the 203 PAS predicted to be regulated by the CU-repeat motifs. 1000 similar-sized sets of poly(A) sites with the same positional preference (distally-located) as the targets of the CU motif were selected and the density of PTBP1 eCLIP reads was computed as described in the Methods section. The mean and standard deviation of position-dependent read density ratios from these randomized data sets are also shown. **(F)** The median ratio of PTBP1-IP to background eCLIP reads over nucleotides 0 to 100 nt downstream of the PAS (position-wise ratios computed as in (E)), for the top 102 (“top”) and bottom 101 (“low”) predicted PTBP1 targets as well as for the background set (“bg”) of distal PAS.

### A novel U-rich motif is associated with 3’ end processing in prostate cancer

Cancer cells, particularly from squamous cell and adenocarcinoma of the lung, express transcripts with shortened 3’ UTRs (Figure 6A, Supplementary Table 4). The negative correlation between the mRNA level expression of CSTF2 and the 3’ UTR length (Figure 6B) led to the suggestion that overexpression of this 3’ end processing factor plays a role in lung cancer [32], Applying KAPAC to 56 matching normal - tumor paired, lung adenocarcinoma samples, we did not find any motifs strongly associated with PAS use changes in this cancer. In particular, we did not recover G/U-rich motifs, as would be expected if CSTF2 were responsible for these changes [32], This was not due to functional compensation by the paralogous CSTF2T, as the expression of CSTF2T was uncorrelated with the 3’ UTR length (Figure 6C). Rather, the CSTF2-specific GU repeat motif had highly variable activity between patients and between poly(A) sites, which did not exhibit a peak immediately downstream of the PAS (Figure 6D), where CSTF2 is known to bind [5], Thus, as in glioblastoma, PAS selection in lung adenocarcinoma likely involves factors other than core 3’ end processing components.

**Figure 6:**
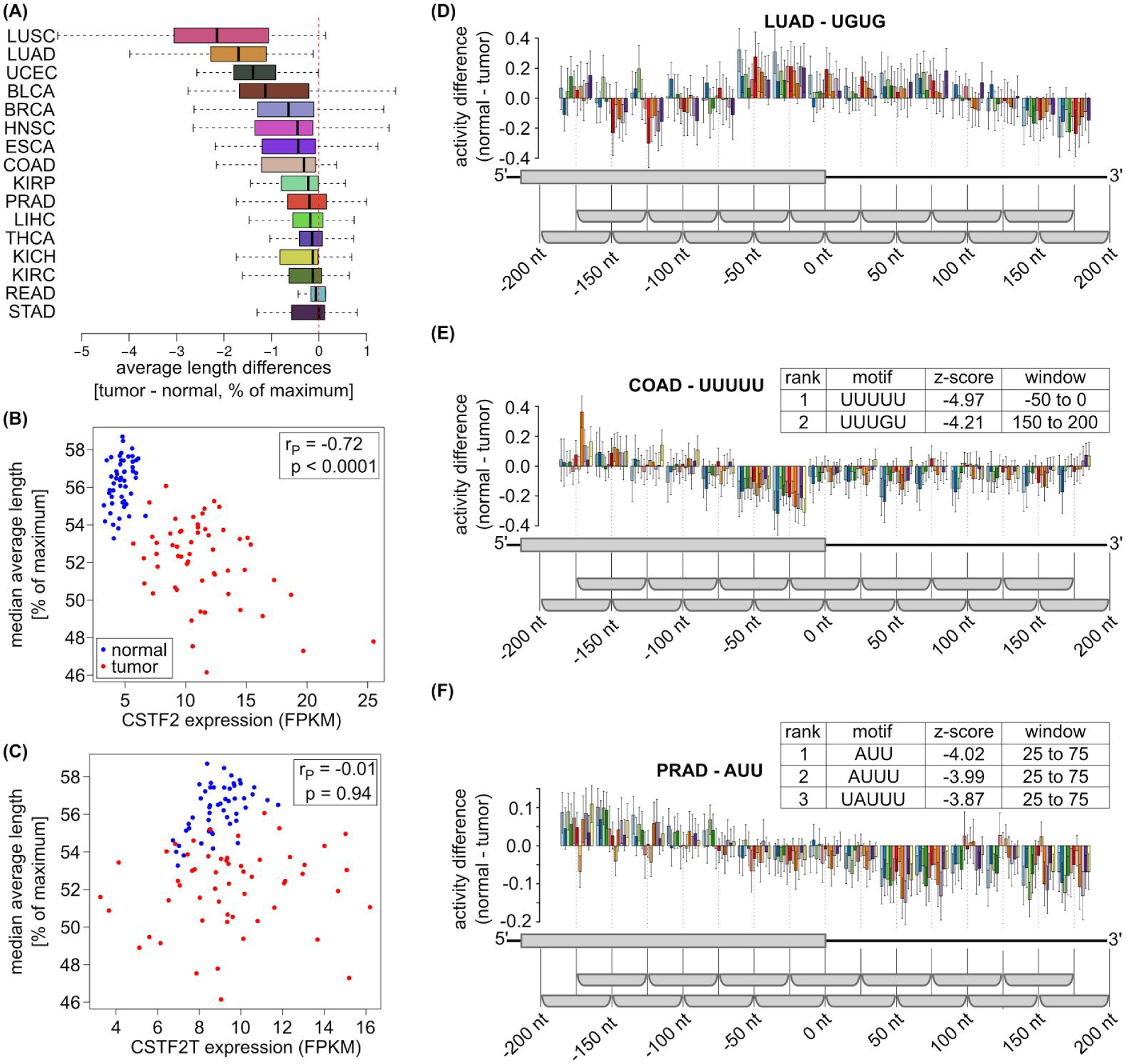
Analysis of TCGA data sets. **(A)** For TCGA data sets with at least 5 matching normal - tumor pairs with high RNA integrity (mTIN > 70), the distributions of patient-wise medians of tumor - normal tissue differences in average terminal exon lengths are shown. Except the adenocarcinoma of the stomach (STAD), the median is negative for all cancers, indicating global shortening of 3’ UTRs in tumors. **(B)** Among 56 matching lung adenocarcinoma (LUAD)-normal tissue pairs (from 51 patients) where global shortening of terminal exons was observed, the CSTF2 expression (in fragments per kilobase per million, FPKM) was negatively correlated (r_P_ = -0.72, p-value = 2.5e-18) with the median of average exon length. **(C)** For the same samples as in (B), no significant correlation (r_P_ = -0.01, p-value = 0.89) between the expression of CSTF2T and the median of average exon length was observed. **(D)** Activity profile of the UGUG CSTF2-binding motif inferred from matched LUAD tumor - normal tissue sample pairs (n = 1054). For visibility, 10 randomly selected sample pairs are shown instead of all 56. **(E, F)** Activity profiles of UUUUU and AUU, the motifs most significantly associated by KAPAC with changes in PAS use in colon adenocarcinoma (COAD, number of PAS n = 1294) **(E)** and prostate adenocarcinoma (PRAD, number of PAS n = 1835) **(F),** respectively (11 tumor - normal tissue sample pairs in both studies).

Exploration of other cancer types for which many paired tumor - normal tissue samples were available revealed that U-rich motifs are more generally significantly associated with changes in PAS use in these conditions (Supplementary Table 3). Most striking was the association of the presence of poly(U) and AUU motifs with an increased PAS use in colon and prostate cancer, respectively (Figure 6E and F). These results indicate that KAPAC can help identify regulators of 3’ end processing in complex tissues environments such as tumors.

## Discussion

Sequencing of RNA 3’ ends has uncovered a complex pattern of PAS and 3’ UTR usage across cell types and conditions, and particularly that the length of 3’ UTRs increases upon cell differentiation and decreases upon proliferation [33, 34], However, the responsible regulators remain to be identified.

The knock-down of most 3’ end processing factors leads to short 3’ UTRs [12], Paradoxically, similar 3’ UTR shortening is also observed in cancers, in spite of a positive correlation between expression of 3’ end processing factors and the proliferative index of cells [3], This may suggest that 3’ end processing factors are not responsible for 3’ UTR processing in cancers, and that other regulators remain to be discovered. However, the possibility remains that 3’ end processing factors, although highly expressed, do not match the increased demand for processing in proliferating cells. Although reduced levels of CFIm 25 have been linked to 3’ UTR shortening and increased tumorigenicity of glioblastoma cells [11], once we applied a threshold on the RNA integrity in the samples to be analyzed, CFIm 25 expression was similar between tumors and normal tissue samples (Supplementary Figure 5). Thus, it seems low apparent CFIm 25 expression is associated with stronger 3’ end bias in read coverage and partial RNA degradation (Supplementary Figure 6). Consistently, our KAPAC analysis of samples with high RNA integrity did not uncover the CFIm 25-specific UGUA motif as significantly explaining the PAS usage changes in glioblastoma compared to normal brain tissue. Of note, in the study of [11] only 60 genes had significantly shortened 3’ UTRs in glioblastoma relative to normal brain, and only 24 of these underwent significant 3’ UTR shortening upon CFIm 25 knock-down in HeLa cells, in spite of 1453 genes being affected by the CFIm 25 knock-down. However, applying KAPAC to 5 normal and 5 glioblastoma tumor samples which showed most separable distributions of terminal exon lengths, we uncovered a pyrimidine motif, likely bound by the PTBP1, as most significantly associated with changes in PAS use in these tumors. Our findings are supported by previous observations that PTBP1 acts antagonistically to CSTF2, repressing PAS usage [35], and that increased PTBP1 expression, as we observed in glioblastoma tumors, promotes proliferation and migration in glioblastoma cell lines [36], Our analysis demonstrates that *de novo*, unbiased motif analysis of tumor data sets with high RNA integrity can reveal specific regulators of PAS usage.

In spite of mounting evidence for the role of CFIm in the regulation of polyadenylation at alternative PAS in terminal exons, its mechanism has remained somewhat unclear. ‘Canonical’ PAS, containing consensus signals for many of the 3’ end processing factors including CFIm, tend to be located distally in 3’ UTRs [5], If core 3’ end processing factors bind to specific PAS and select them for processing, reducing the concentration of 3’ end processing factors should increase the stringency of PAS selection. Yet the siRNA-mediated knock-down of CFIm leads to increased processing at proximal sites, and not to preferential processing of the ‘high-affinity’, distal PAS. Here we have found that CFIm indeed promotes the usage of distal PAS to which it binds, while CFIm binding motifs are depleted at both the proximal and the distal PAS of terminal exons whose processing is insensitive to the level of CFIm. Therefore, the decreased processing of distal PAS upon CFIm knock-down is not explained by a decreased ‘affinity’ of these sites. A model that remains compatible with the observed pattern of 3’ end processing is the so-called ‘kinetic’ model, whereby reducing the rate of processing at a distal, canonical site when the regulator is limiting, leaves sufficient time for the processing of a suboptimal proximal site. Kinetic aspects of pre-mRNA processing have started to be investigated in cell lines that express slow and fast-transcribing RNA polymerase II (RNAPII) [37], Analyzing RNA-seq data from these cells, we found that terminal exons that respond to CFIm knock-down in our data, underwent more pronounced shortening in cells expressing the slow polymerase (Supplementary Figure 7), in agreement with the kinetic model. Nevertheless, this effect was also apparent for exons in which proximal and distal poly(A) sites were located far apart, it was not limited to CFIm targets. Furthermore, the changes in 3’ UTR length in a sample from the fast RNAPII-expressing cell line were surprisingly similar to the changes we observed for the slow polymerase. Thus, current data do not provide unequivocal support to the kinetic model underlying the relative increase in processing of proximal PAS upon CFIm knock-down.

Generalized linear models have been widely used to uncover transcriptional regulators that implement gene expression programs in specific cell types [38, 39], Similar approaches have not been applied to 3’ end processing, possibly because the genome-wide mapping of 3’ end processing sites has been lagging behind the mapping of transcription start sites. Here we demonstrate that the modeling of PAS usage in terms of motifs in the vicinity of PAS can reveal global regulators, while the reconstructed position-dependent activity of their corresponding motifs provides insights into their mechanisms. Interestingly, some of the proteins that we touched upon in our study are splicing factors. This underscores a general coupling between splicing and polyadenylation that has been long surmised (e.g. [17]), and for which evidence has started to emerge [40], Interestingly, the activities of splicing factors on poly(A) site choice paralleled the activities of these factors on splice site selection. Specifically, we found that both HNRNPC, which functions as an ‘RNA nucleosome’ in packing RNA and masking decoy splice sites [24], and PTBP1 which has repressive activity on exon inclusion [41], repress the processing of the PAS to which they bind. This unexpected concordance in activities suggests that other splicing factors simultaneously modulating 3’ end processing are to be uncovered. Splicing is strongly perturbed in cancers [42], and the role of splicing factors in the extensive change of the polyadenylation landscape remains to be defined.

Sequencing of RNA 3’ ends has greatly facilitated the study of 3’ end processing dynamics. However, such data remain relatively uncommon, and many large-scale projects have already generated a wealth of RNA sequencing data that could, in principle, be mined to uncover regulators of CPA. We found a previously proposed method for inferring the relative use of alternative PAS from RNA-seq data, DaPars [11], to have limited reproducibility, possibly because biases in read coverage along RNAs are difficult to model. To overcome these limitations, we developed PAQR, which makes use of a large catalog of PAS to segment the 3’ UTRs and infer the relative use of PAS from RNA-seq data. We show that PAQR enables a more reproducible as well as accurate inference of motif activities in PAS choice compared to DaPars. PAQR strongly broadens the domain of applicability of KAPAC to include RNA sequencing data sets that have been obtained in a wide range of systems, as we have illustrated in our study of TCGA data. As single-cell transcriptome analyses currently employ protocols designed to capture RNA 3’ ends, it will be especially interesting to apply our methods to single-cell sequencing data.

## Conclusions

In this study, we developed PAQR, a robust computational method for inferring relative poly(A) site use in terminal exons from RNA sequencing data and KAPAC, an approach to infer sequence motifs that are associated with the processing of poly(A) sites in specific samples. We demonstrate that these methods help uncover regulators of polyadenylation in cancers and also shed light on their mechanism of action. Our study further underscores the importance of assessing the quality of samples used for high-throughput analyses, as this can have substantial impact on the estimates of gene expression.

## Methods

### Datasets

#### A-seq2 samples

3’ end sequencing data from HeLa cells that were treated with either a control siRNA or siRNAs targeting the CFIm 25 and the CFIm 68 transcripts were generated as follows. HeLa cells were cultured in DMEM (# D5671, Sigma Aldrich) supplemented with L Glutamine (#25030081, ThermoFisher Scientific) and 10% fetal bovine serum (#7524, Sigma-Aldrich). For siRNA treatment, cells were seeded in 6 well polystyrene-coated microplates and cultured to reach a confluence of ∼50%. Subsequently, the cells were separately transfected with 150 picomoles of siRNA, either control (sense strand sequence - 5’ AGG UAG UGU AAU CGC CUU GTT 3’), or directed against CFIm 25 (sense strand sequence - 5’ GGU CAU UGA CGA UUG CAU UTT 3’) or against CFIm 68 (sense strand sequence - 5’ GAC CGA GAU UAC AUG GAU ATT 3’), with Lipofectamine RNAiMAX reagent (#13778030, ThermoFisher Scientific). All siRNAs were obtained from Microsynth AG and had dTdT overhangs. The cells were incubated with the siRNA Lipofectamine RNAiMax mix for at least 48 hours before cells were lysed. Cell lysis and polyadenylated RNA selection was performed according to the manufacturer’s protocol (Dynabeads™ mRNA DIRECT™ Purification Kit, #61011, Thermo Scientific). Polyadenylated RNA was subsequently processed and libraries were prepared for sequencing on the Illumina HiSeq 2500 platform as described earlier [19], Sequencing files were processed according to Martin et al. [43] but without using the random 4-mer at the start of the sequence to remove duplicates. A-seq2 3’ end processing data from control and si-HNRNPC-treated cells was obtained from a prior study [19].

### 3’ end sequencing data pertaining to PCBP1

3’ end sequencing data from control and si-PCPB1-treated cells were downloaded from SRA (accession: SRP022151) and converted to fastq format. Reverse complemented and duplicate-collapsed reads were then mapped to the human genome with segemehl version 0.1.7 [44], We did not use STAR for these data set because these libraries, generated by DRS (direct RNA sequencing) had a high fraction of short reads that STAR did not map. From uniquely mapped reads for which at least the last 4 nucleotides at the 3’ end perfectly matched to the reference, the first position downstream of the 3’ end of the alignment was considered as cleavage site and used for quantification of PAS use.

### RNA-seq data from The Cancer Genome Atlas

BAM files for matching normal and tumor RNA-seq samples listed in Supplementary Table 5 were obtained from the Genomic Data Commons (GDC) Data Portal [45] along with gene expression values normalized with HTSeq and reported in fragments per kilobase per million (FPKM).

### Other RNA-seq data sets

Publicly available raw sequencing data were obtained from NCBI’s Gene Expression Omnibus (GEO) [46] for the studies of CFIm 25 knock-down in HeLa cells [11] (accession number GSE42420), HNRNPC knock-down in HEK293 cells [29] (GSE56010), PTBP1/2 knock-down in HEK293 cells [30] (GSE69656) and for HEK293 cells expressing mutated versions of POLR2A that have overall different rates of RNAPII transcription elongation [37] (GSE63375).

### PTBP1 CLIP data

PTBP1-eCLIP data generated by the ENCODE consortium [31] was obtained from the ENCODE Data Coordination Center [47] (accession numbers for the IP and control samples from K562 cells ENCSR981WKN and ENCSR445FZX, and from HepG2 cells ENCSR384KAN and ENCSR438NCK).

### Processing of the sequencing data

Raw reads obtained from RNA-seq experiments were mapped according to the RNA-seq pipeline for long RNAs provided by the ENCODE Data Coordinating Center [48] using the GENCODE version 24 human gene annotation. Raw reads from the study conducted by Gueroussov et al. [30] were additionally subjected to 3’ adapter trimming with cutadapt, version 1.14 [49] prior to mapping. Raw reads from eCLIP experiments carried out by the ENCODE consortium for the PTBP1 were first trimmed with cutadapt version 1.9.1 [49], both at the 5’ and at the 3’ ends to remove adapters. A second round of trimming guaranteed that no double ligation events were further processed. The reads were then mapped to the genome with STAR, version 2.5.2a [50], Detection and collapsing of PCR duplicates was done with a custom python script similar to that described by van Nostrand et al. [27], BAM files corresponding to biological replicates were then merged.

## PAQR

### Inputs

PAQR requires an alignment file in BAM-format and a file with all poly(A) sites mapped on the genome, in BED-format. The assessment of RNA integrity (see below) also requires the transcript annotation of the genome, in BED12-format.

### Poly(A)sites

PAQR quantifies the relative use of poly(A) sites in individual terminal exons. We started from the entire set of poly(A) sites in the PolyAsite resource [19], but this set can be exchanged or updated, and should be provided as a BED-file to the tool. We converted the coordinates of the poly(A) sites to the latest human genome assembly version, GRCh38, with liftOver [51], Terminal exons with more than one poly(A) site (terminal exons with tandem poly(A) sites, TETPS) and not overlapping with other annotated transcripts on the same strand were identified based on version 24 of the GENCODE [52] annotation of the genome. When analyzing RNA-seq data that was generated with an unstranded protocol, PAQR does not quantify poly(A) site usage in terminal exons that overlap with annotated transcripts on the opposite strand.

### Quantification of PAS usage

The main steps of the PAQR analysis are as follows: first, the quality of the input RNA sequencing data is assessed, to exclude samples with evidence of excessive RNA degradation. Samples that satisfy a minimum quality threshold are then processed to quantify the read coverage per base across all TETPS and poly(A) sites with sufficient evidence of being processed are identified. These are called ‘used’ poly(A) sites (or uPAS). Finally, the relative use of the uPAS is calculated.

### Assessment of sample integrity

The integrity of RNA samples is usually assessed based on a fragment analyzer profile [53], Alternatively, a post hoc method, applicable to all RNA sequencing data sets, quantifies the uniformity of read coverage along transcript bodies in terms of a ‘transcript integrity number’ (TIN) [54], We implemented this approach in PAQR, calculating TIN values for all transcripts containing TETPS. For the analysis of TCGA samples and of RNA-seq samples from cells with different RNAPII transcription speeds, we only processed samples with a median TIN value of at least 70, as recommended in the initial publication [54].

### RNA-seq read coverage profiles

For each sample, nucleotide-wise read coverage profiles along all TETPS were calculated based on read-to-genome alignments (obtained as described above). In processing paired-end sequencing data, PAQR ensured unique counting of reads where the two mates overlap. When the data was generated with an unstranded protocol, all reads that mapped to the locus of a specific TETPS were assumed to originate from that exon. The locus of each TETPS was extended by 200 nt at the 3’ end, to ensure inclusion of the most distal poly(A) sites (see below). To accurately quantify the usage of the most proximal PAS, when poly(A) sites were located within 250 nt from the start of the terminal exon, the coverage profile was first extended upstream of the PAS based on the reads that mapped to the upstream exon(s). Specifically, from the spliced reads, PAQR identified the upstream exon with most spliced reads into the TETPS and computed its coverage. When the spliced reads that covered the 5’ end of the TETPS provided evidence for multiple splice events, the most supported exons located even further upstream were also included (Supplementary Figure 8).

### Identification of the most distal poly(A) sites

From the read coverage profiles, PAQR attempted to identify the poly(A) sites that show evidence of processing in individual samples as follows. First, to circumvent the issue of incomplete or incorrect annotations of PAS in transcript databases, PAQR identified the most distal PAS in each terminal exon that had evidence of being used in the samples of interest. Thus, alignment files were concatenated to compute a joint read coverage profile from all samples of the study. Then, the distal PAS was identified as the 3’-most PAS in the TETPS for which: 1. The mean coverage in the 200 nt region downstream of the PAS was lower than the mean coverage in a region twice the read length (to improve the estimation of coverage, as it tends to decrease towards the poly(A) site) upstream of the poly(A) site, and 2. The mean coverage in the 200 nt region downstream of the PAS was at most 10 % of the mean coverage from the region at the exon start (the region within one read length from the exon start) (Supplementary Figure 9). For samples from TCGA, where read length varied, we have used the maximum read length in the data for each cancer type. After the distal PAS was identified, PAQR considered for the relative quantification of PAS usage only those TETPS with at least one additional PAS internal to the TETPS and with a mean raw read coverage computed over the region between the exon start and distal PAS of more than five.

### Identification of used poly(A) sites

PAQR infers the uPAS recursively, at each step identifying the PAS that allows the best segmentation of a particular genomic region into upstream and downstream regions of distinct coverage across all replicates of a given condition ( Figure 3). Initially, the genomic region is the entire TETPS, and at subsequent steps genomic regions are defined by previous segmentation steps. Given a genomic region and annotated PAS within it, every PAS is evaluated as follows. The mean squared error (MSE) in read coverage relative to the mean is calculated separately for the segments upstream (MSE_u_) and the downstream (MSE_d_) of each PAS for which the mean coverage in the downstream is lower than the mean coverage in the upstream region. A minimum length of 100 nt is required for each segment, otherwise the candidate PAS is not considered further. The sum of MSE in the upstream and downstream segments is compared with the MSE computed for the entire region (MSE_t_). If (MSE_u_+MSE_d_)/MSE_t_ ≤ 0.5 (see also below), the PAS is considered ‘candidate used’ in the corresponding sample. When the data set contains at least 2 replicates for a given condition, PAQR further enforces the consistency of uPAS selection in replicate samples by requiring that the PAS is considered used in at least 2 of the replicates and furthermore, for all PAS with evidence of being used in a current genomic region, the one with the smallest median MSE ratio computed over samples that support the usage of the site is chosen in a given step of the segmentation. The segmentation continues until no more PAS has sufficient evidence of being used. If the data consists of a single sample, the segmentation is done based on the smallest MSE at each step.

To further minimize incorrect segmentations due to PAS that are used in the samples of interest but not part of the input set, an additional check is carried out for each TETPS in each sample, to ensure that applying the segmentation procedure considering all positions in the TETPS rather than the annotated PAS recovers positions that fall within at most 200 nt upstream of the uPAS identified in previous steps for each individual sample (Supplementary Figure 10). If this is not the case, the data for the TETPS from the corresponding sample is excluded from further analysis.

### Treatment of closely spaced poly (A) sites

Occasionally, distinct PAS occur very close to each other. While 3’ end sequencing may allow their independent quantification, the RNA-seq data does not have the resolution to distinguish between closely spaced PAS. Therefore, in the steps described above, closely spaced (within 200 nt of each other) PAS are handled first, to identify one site of the cluster that provides the best segmentation point. Only this site is then compared with the more distantly spaced PAS.

### Relative usage and library size normalized expression calculation

Once used poly(A) sites have been identified, library size-normalized expression levels and relative usage within individual terminal exons are calculated. Taking a single exon in a single sample, the following steps are performed: the mean coverage of the longest 3’ UTR is inferred from the region starting at the most distal poly(A) site and extending upstream up to the next poly(A) site or to the exon start. Mean coverage values are similarly calculated in regions between consecutive poly(A) sites and then the coverage of individual 3’ UTR is determined by subtracting from the mean coverage in the terminal region of that 3’ UTR the mean coverage in the immediately downstream region. As some of the poly(A) sites are not identified in all samples, their usage in the samples with insufficient evidence is calculated as for all other sites, but setting the usage to 0 in cases in which the upstream coverage in the specific sample was lower than the downstream coverage. The resulting values are taken as raw estimates of usage of individual poly(A) sites and usage relative to the total from poly(A) sites in a given terminal exon are obtained.

To obtain library size normalized expression counts, raw expressions from all quantified sites of a given sample are summed up. Each raw count is divided by the summed counts (i.e. the library size) and multiplied by 10^6^, resulting in expression estimates as reads per million (RPM).

### PAQR modules

PAQR is composed of 3 modules: (1) A script to infer transcript integrity values based on the method described in a previous study [54], The script builds on the published software which is distributed as part of the Python RSeQC package version 2.6.4 [55], (2) A script to create the coverage profiles for all considered terminal exons. This script relies on the HTSeq package version 0.6.1 [56] and (3) a script to obtain the relative usage together with the estimated expression of poly(A) sites with sufficient evidence of usage.

All scripts, intermediate steps, and analysis of the TCGA data sets were executed as workflows created with snakemake version 3.13.0 [57].

### KAPAC

KAPAC, standing for K-mer Activity on Polyadenylation Site Choice, aims to identify k-mers that can explain the change in PAS usage observed across samples. For this, we model the relative change in PAS-usage within terminal exons (with respect to the mean across samples) as a linear function of the occurrence of a specific k-mer and the unknown “activity” of this k-mers. Note that by modeling the relative usage of PAS within individual terminal exons we will capture only the changes that are due to alternative polyadenylation and not those that are due to overall changes in transcription rate or to alternative splicing. We are considering k-mers of a length from 3 to 6 nucleotides in order to match the expected length of RBP binding sites [28].

KAPAC attempts to explain the change in the relative use of a given PAS in terms of the motifs (k-mers) that occur in its vicinity, each occurrence of a k-mer contributing a multiplicative constant to the site use. Thus, we write the number of reads observed from PAS *i* in sample s as *R_i,s_* = α* *exp*(*N_i,k_*\**A_k,s_*), where *N_i,k_* is the count of k-mer *k* around PAS *i*, *A_k,s_* the activity of the k-mer in sample *s*, which determines how much the k-mer contributes to the PAS use, and *α* is the overall level of transcription at the corresponding locus. Then, for poly(A) sites in the same terminal exon we can write their log relative use *log*_2_ (*U_i,s_*) as function of the number of k-mer counts found in a defined window at a specific distance from the site *i* and the activity of these k-mers: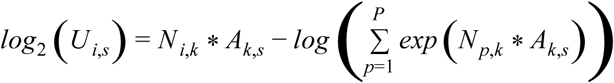 (see Supplementary methods for a detailed derivation). By fitting the relative use of poly(A) sites to the observed number of motifs around them, we can obtain the activities *A_k,s_* for each k-mer *k* in each sample s and calculate mean activity difference z-scores across treatment versus control pairs of samples (see Figure 1 and Supplementary methods).

### Parameters used for KAPAC analysis of 3’ end sequencing data

We considered terminal exons with multiple poly(A) sites within protein coding transcripts (hg38, GENCODE version 24) whose expression, inferred as previously described [19], was at least 1 RPM in at least one of the investigated samples. To ensure that the position-dependent motif activities could be correctly assigned, exons containing expressed PAS that were closer than 400 nt from other PAS were excluded from the analysis, as we applied KAPAC to regions +/-200 nt around poly(A) sites. We randomized the associations of changes in poly(A) site use with k-mer counts 100 times in order to calculate p-values for mean activity difference z-scores (see Supplementary methods).

### Parameters used for KAPAC analysis of RNA-seq data

All KAPAC analyses for RNA-seq data sets considered terminal exons with at least 2 PAS of any transcripts from the GENCODE version 24 annotation of the human genome. Filtering of the closely-spaced PAS, activity inference and randomization tests were done similar to the processing of 3’ end sequencing libraries. No TPM cutoff was applied as the used PAS are already determined by PAQR.

### Average terminal exon length

An average terminal exon length can be calculated over all transcripts expressing a variant of that terminal exon as 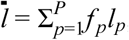, where *f_p_* is the relative frequency of use of PAS p in the terminal exon and *l_p_* is the length of the terminal exon when PAS p is used for CPA. To compare terminal exons with different maximum lengths, we further normalize the average exon length to the maximum and express this normalized value percentually. Thus, when the most distal site is exclusively used the average terminal exon length is 100, while when a very proximal site is used exclusively, the average terminal exon length will be close to 0 (Supplementary Figure 11).

### Average length difference

The difference in average length of a terminal exon between two samples is obtained by subtracting the average length inferred from one sample from the average length inferred from the second sample. 3’ UTR shortening is reflected in negative average length differences, while 3’ UTR lengthening will lead to positive differences.

### Definition of the best MSE ratio threshold

Two studies of HNRNPC yielded 3’ end sequencing [19] and RNA sequencing [29] data of control and si-HNRNPC-treated cells. We used these data to define a PAQR parameter (the threshold MSE ratio) such as to maximize the reproducibility of the results from the two studies. MSE ratio values ranging from 0.2 to 1.0 were tested, Supplementary Figure 12. Relative use of PAS we calculated based on the A-seq2 data sets as described before [19], The RNA-seq data was processed to infer PAS use with different MSE cutoffs, and the calculate average terminal exon lengths for individual exons in individual samples and also differences in average exon lengths between samples. For the comparison of the RNA-seq based PAS quantifications with those from A-seq2, we considered both the overall number of terminal exons quantified in replicate data sets as well as the correlation of average length differences. As shown in Supplementary Figure 12 stringent (low) cutoff in MSE leads to few exons being quantified with high reproducibility, but the number of quantified exons has a peak relative to the MSE. At a threshold of 0.5 on MSE we are able to quantify the largest number of exons with relatively good reproducibility, and we therefore applied this value for all our subsequent applications of PAQR.

### Selection of normal - tumor sample pairs for analysis of 3’ UTR shortening

For the analysis of motifs associated with 3’ UTR length changes in cancers, we computed the distribution of 3’ UTR length differences in matched tumor-normal samples. We carried out hierarchical clustering of vectors of 3’ UTR length changes for each cancer type separately (using Manhattan distance and complete linkage). We then identified the subcluster in which the median change in 3’ UTR length was negative for all samples and that also contained the sample where the median change over all transcripts had the median value over all samples. Samples from these clusters were further analyzed with KAPAC.

### Selection of normal - tumor pairs from GBM data

From the six normal tissue sample that had a median transcript integrity number > 0.7, five had similar average exon length distributions (all of them being among the samples with the highest median average length). We used these five normal tissue samples and selected five primary tumor samples with similarly high TIN and the lowest median average exon length. We then generated random pairs of normal-tumor tissue samples and analyzed them similarly to paired samples from other cancers.

### eCLIP data analysis

We predicted targets of the CU-repeat motif as described in the Supplementary methods and obtained a total of 203 targets. We either used the entire set or divided the set into the top half and bottom half of targets. For each poly(A) site from a given set, the read coverage profiles of the 400 nt region centered on the poly(A) site were constructed from both the protein-specific immunoprecipitation (IP) experiment and the related size-matched control. At every position, we computed the ratio of the library size normalized read coverage (RPM) in the IP and in the background sample (using a pseudo-count of 0.1 RPM) and then average these ratios position-wise across all poly(A) sites from a given set, considering any poly(A) site with at least a single read support in either of both experiments. For comparison, we carried out the same analysis for 1000 random sets of poly(A) sites with the same size as the real set, and then inferred the mean and standard deviation of the mean read ratios at each position.

### Motif profiles

Motif profiles were generated by extracting the genomic sequences (from the GRCh38 version of the human genome assembly) around poly(A) sites from a given set, scanning these sequences and tabulating the start positions where the motif occurred. The range of motif occurrence variation at a given position was calculated as the standard deviation of the mean, assuming a binomial distribution with the probability of success given by the empirical frequency (smoothened over 7 nucleotides centered on the position of interest) and the number of trials given by the number of poly(A) sites in the set.

### Selection of CFIm-sensitive and insensitive terminal exons

For terminal exons with exactly two quantified poly(A) sites that were expressed with at least 3 RPM in all samples (1776 terminal exons) we calculated the proximal-distal ratio. Next, we calculated the average (between replicates) log10 fold-change (in knock-down relative to control) in proximal-distal ratio. The 750 terminal exons with the largest average Iog10 fold-change in the CFIm 25 and CFIm 68 knock-down experiments were selected as CFIm sensitive, while the 750 with an average Iog10 fold-change closest to zero were considered insensitive.

## List of abbreviations

TGCA cancer cohort abbreviations used in the manuscript correspond to the following full names:

BCLA: Bladder Urothelial Carcinoma
BRCA: Breast Invasive Carcinoma
COAD: Colon Adenocarcinoma
ESCA: Esophageal Carcinoma
GBM: Glioblastoma Multiforme
HNSC: Head and Neck Squamous Cell Carcinoma
KICH: Kidney Chromophobe
KIRC: Kidney Renal Clear Cell Carcinoma
KIRP: Kidney Renal Papillary Cell Carcinoma
LIHC: Liver Hepatocellular Carcinoma
LUAD: Lung Adenocarcinoma
LUSC: Lung Squamous Cell Carcinoma
PRAD: Prostate Adenocarcinoma
READ: Rectum Adenocarcinoma
STAD: Stomach Adenocarcinoma
THCA: Thyroid Carcinoma
UCEC: Uterine Corpus Endometrial Carcinoma

## Declarations

### Availability of data and materials

3’ end sequencing data from HeLa cells treated with control siRNA or siRNAs directed against CFIm 25 and CFIm 68 and generated with the A-seq2 protocol [43] have been submitted to the NCBI Sequence Read Archive (SRA) [58] and are available under accession number SRP115462. A-seq2 data pertaining to HNRNPC were obtained from SRA under accession number SRP065825. Direct RNA sequencing data from the PCBP1 study of Ji et al. [23] were obtained from SRA with accession number SRP022151. RNA sequencing data from the studies involving CFIm 25 knock-down [11], HNRNPC knock-down [29], PTBP1/2 knock-down [30] and RNAPII with altered elongation rate [37] were obtained from GEO [46], with accession numbers GSE42420, GSE56010, GSE69656, and GSE63375, respectively. Data from the eCLIP study of PTBP1 was obtained from the ENCODE Data Coordination Center [47], having the following accession numbers: ENCSR981WKN, ENCSR445FZX, ENCSR384KAN and ENCSR438NCK. The TCGA data was obtained from the GDC Portal [45], following permission.

## Acknowledgements

We are grateful to the specimen donors and to the research groups that were part of the TCGA research network for making these data available. We would like to thank Florian Geier for fruitful discussions and sharing R code for regression models. Also, we would like to thank the sciCORE team for their maintenance of the HPC facility at the University Basel and John Baumgartner for his R implementation of Iwanthue (https://github.eom/johnbaums/hues/blob/master/R/iwanthue.R).

## Authors’ contributions

Andreas J. Gruber developed KAPAC, Ralf Schmidt developed PAQR, Souvik Ghosh and Georges Martin generated the 3’ end sequencing data in HeLa cells, Andreas R. Gruber contributed to the analysis of 3’ end sequencing data sets with KAPAC and Erik van Nimwegen contributed to the KAPAC model. AJG and RS analyzed the 3’ end and RNA sequencing data sets. Mihaela Zavolan contributed to model development and analyses. AJG, RS, MZ wrote the manuscript with help from all authors.

## Ethics approval and consent to participate

Authorization to use RNA-seq data from patient samples, which is obtained by the TCGA Research Network, has been granted.

